# Identification of a pharmacokinetic interaction between teicoplanin and sulfo-butyl ether-β-cyclodextrin, an excipient in the intravenous posaconazole

**DOI:** 10.64898/2026.02.17.706257

**Authors:** Yoshimasa Adachi, Mitsuhiro Sugimoto, Yusei Yamada, Junya Kanda, Atsushi Yonezawa, Takeo Yamagiwa, Yuta Hanyu, Mizuki Watanabe, Yasuyuki Arai, Chisaki Mizumoto, Toshio Kitawaki, Tadakazu Kondo, Kouhei Yamashita, Natsuki Imayoshi, Yuki Shigetsura, Yurie Katsube, Keiko Ikuta, Daiki Hira, Ryuji Ikeda, Akifumi Takaori-Kondo, Shunsaku Nakagawa, Tomohiro Terada

**Author notes:** Address correspondence to Shunsaku Nakagawa. Department of Clinical Pharmacology and Therapeutics Kyoto University Hospital, 54 Kawahara-cho, Shogoin, Sakyo-ku, Kyoto 606-8507, Japan.

## Abstract

Patients undergoing hematopoietic stem cell transplantation (HSCT) often receive multiple antibiotics and antifungals concurrently, making it crucial to understand their potential pharmacokinetic interactions of these agents. We report here an interaction between the glycopeptide antibiotic teicoplanin (TEIC) and sulfo-butyl ether-β-cyclodextrin (SBECD), a solubilizing excipient in the intravenous formulation of posaconazole (PSCZ). We performed a single-center retrospective analysis of patients who underwent HSCT and received oral and intravenous PSCZ during TEIC therapy. The associations between PSCZ administration and TEIC concentration-to-dose (C/D) ratios were evaluated using linear mixed-effects models. We also examined the effects of intravenous PSCZ and SBECD on TEIC pharmacokinetics in rats by assessing the area under the concentration–time curve (AUC) and urinary excretion of total TEIC and its components. In addition, molecular docking and in vitro protein-binding assays were conducted to investigate the interaction between TEIC and SBECD. In patients who underwent HSCT, TEIC C/D ratio was significantly lower following intravenous PSCZ administration than without administration. In contrast, the effect of oral PSCZ administration relative to non-administration was not statistically significant. In rats, intravenous PSCZ and SBECD decreased the AUC of TEIC and increased urinary excretion, particularly in the A_2_ group. Docking simulations indicated that the hydrophobic side chain of TEIC A_2-2_ fit within the SBECD cavity, and in vitro assays confirmed SBECD concentration-dependent increases in TEIC unbound fractions. The co-administration of intravenous PSCZ containing SBECD may reduce TEIC protein binding, thereby enhancing renal excretion.

## Introduction

Patients who undergo hematopoietic stem cell transplantation (HSCT) are susceptible to bacterial and fungal infections owing to immunosuppression caused by conditioning chemotherapy before transplantation and subsequent immunosuppressive therapy. Consequently, multiple antibiotics and antifungal agents are often administered concomitantly to these patients for the prophylaxis and treatment of infections.

Teicoplanin (TEIC), a glycopeptide antibiotic, is one of the agents frequently used in this population. Along with vancomycin, TEIC is widely used in the treatment of methicillin-resistant *Staphylococcus aureus* infections, and is administered empirically when gram-positive bacterial infections are suspected. TEIC undergoes minimal metabolism and is eliminated primarily through the kidneys. Compared to vancomycin, TEIC exhibits a higher protein-binding rate, a larger volume of distribution, and a substantially longer half-life of approximately 83–168 h (1, 2). Target trough concentrations required for therapeutic efficacy have been established based on pharmacokinetic-pharmacodynamic analyses (3, 4); thus, maintaining appropriate plasma concentrations is critical for successful treatment. To this end, fluctuations in TEIC exposure may result in treatment failure, highlighting the need to fully elucidate inter- and intra-individual factors that contribute to pharmacokinetic variability.

Azole antifungal agents are commonly administered to patients undergoing HSCT to prevent fungal infections. Posaconazole (PSCZ), an azole antifungal agent, is known for its potent antifungal activity and inhibition of the drug-metabolizing enzyme CYP3A and efflux transporter P-glycoprotein in the liver and small intestine (5, 6).

Therefore, co-administration of PSCZ with drugs that are substrates of these enzymes or transporter may reduce the clearance of the co-administered drugs, potentially leading to drug–drug interactions. In contrast, PSCZ is unlikely to significantly affect the pharmacokinetics of renally excreted drugs. Patients undergoing HSCT often receive multiple antibiotics and antifungals concurrently; however, data on the pharmacokinetic interactions among these agents in this population remain limited.

Therefore, the aim of this study was to evaluate the effect of intravenous (i.v.) PSCZ co-administration on the plasma concentration of TEIC in patients undergoing HSCT. We assessed the reproducibility of this phenomenon using an animal model, and further identified sulfo-butyl ether-β-cyclodextrin (SBECD), a pharmaceutical excipient contained in i.v. PSCZ, as a potential contributor to the reduction in plasma TEIC concentrations. Moreover, we predicted the formation of an inclusion complex of SBECD with TEIC, and evaluated the effect of SBECD on TEIC protein binding.

## Materials and Methods

### Effects of oral and intravenous PSCZ administration on TEIC plasma concentrations in patients who underwent HSCT

We conducted a single-center retrospective analysis of patients who underwent HSCT and received the tablet formulation (oral) and injectable formulation of PSCZ (i.v. PSCZ) during TEIC treatment between April 2020 and January 2022 at the Department of Hematology, Kyoto University Hospital. Eligible patients were required to have available plasma TEIC concentration data under three conditions: without PSCZ administration, following oral PSCZ administration, and following i.v. PSCZ administration. This study was conducted in accordance with the Declaration of Helsinki and its amendments and was approved by the Ethics Committee of Kyoto University Graduate School and Faculty of Medicine (Approval No. R2470).

We extracted all available data on trough TEIC plasma concentrations from the hospital electronic medical database. Plasma TEIC concentrations were measured in the hospital’s central laboratory by latex immunoturbidimetry using the Cobas 6000 system. TEIC plasma concentration data were included if they met the following criteria: (i) obtained at least 3 days after maintaining a constant dosage following TEIC loading, (ii) obtained at least 3 days after dosage adjustment, or (iii) obtained at least 3 days after switching the PSCZ formulation. We also collected data on sex and age, which are generally assumed to influence drug pharmacokinetics, as well as serum albumin levels and estimated glomerular filtration rate (eGFR), which are known to affect the pharmacokinetics of TEIC (7).

To evaluate the association between PSCZ administration and changes in TEIC pharmacokinetics, the effects of PSCZ on the concentration-to-dose (C/D) ratio of TEIC were estimated using a linear mixed-effects model:

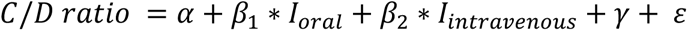

where *α* is the estimate of the intercept, *β* is the estimate of the fixed effect, *I*_oral_ and *I*_intravenous_ are the indicator variables, *γ* is the estimate of the random effect for the intercept (individual-level residual with mean zero and variance σ^2^_γ_), and *ε* is the estimated residual error (with mean zero and variance σ^2^_ε_). *I_oral_* = 1 if oral PSCZ was co-administered, and 0 otherwise; *I_intravenous_* = 1 if i.v. PSCZ was co-administered, and 0 otherwise. The effects of PSCZ were further adjusted for age, sex, serum albumin level, and eGFR, which were incorporated as independent covariates with fixed effects. Analyses were performed using JMP software version 18 (SAS Institute Inc., Cary, NC, U.S.A.)

### Pharmacokinetics of TEIC in rats

Eight-week-old male Wistar/ST rats were purchased from the SLC Animal Research Laboratories (Shizuoka, Japan) and maintained in accordance with the guidelines for Animal Experiments of Kyoto University, Kyoto, Japan. The study was approved by the Animal Research Committee of the Graduate School of Medicine, Kyoto University, Kyoto, Japan (permit numbers: Medkyo 22119 and 23107). The rats were housed in a specific pathogen-free facility, maintained in a temperature-controlled environment with a 12-h light/dark cycle, and fed a standard diet and water ad libitum. For the administration of TEIC and PSCZ to rats, the commercially available i.v. formulations, Teicoplanin for Intravenous Infusion 200 mg (Towa) (Towa Pharmaceutical Co., Ltd. Osaka, Japan) and Noxafil® for Intravenous Infusion 300 mg (MSD K.K., Tokyo, Japan) were used. SBECD was purchased from ChemScene (NJ, USA). All other reagents and solvents were of analytical or liquid chromatography-mass spectrometry grade.

The rats were anesthetized with a combination of medetomidine, midazolam, and butorphanol, and catheters were inserted into the left femoral vein and right femoral artery using polyethylene tubing (SP-31; Natsume Seisakusho, Tokyo, Japan). Saline, PSCZ (40 mg/kg PSCZ plus 890.67 mg/kg SBECD), or SBECD alone (890.67 mg/kg) was administered through the femoral vein. These doses were set with reference to the human equivalent dose in rats, where the dose per body weight is approximately six times the human dose (8). Thirty minutes later, TEIC (20 mg/kg) was injected into the femoral vein. Six rats per group were used in the experiment. PSCZ is highly hydrophobic, making it difficult to dissolve in a solvent suitable for i.v. administration without SBECD. Therefore, the PSCZ-alone group was not included in this study.

Blood samples were collected from the femoral artery at the end of TEIC administration and 5, 15, 30, 60, 90, and 120 min thereafter. Plasma was separated by centrifugation.

Urine was collected from the bladder into a polyethylene tube for 120 min after TEIC administration. Plasma and urine samples were deproteinized by adding four volumes of methanol, followed by centrifugation at 10,000 × *g* for 5 min. A 50 μL aliquot of the supernatant was mixed with 200 μL of methanol. Urine samples were further diluted five-fold with methanol. Lastly, all samples were filtered through a 0.45 µm membrane filter prior to analysis.

TEIC was quantified using a triple-quadrupole mass spectrometry system (LCMS-8040; Shimadzu, Japan) equipped with an electrospray ionization source. The mobile phases were 0.1% formic acid in water (phase A) and 0.1% formic acid in acetonitrile (phase B). The gradient elution program was as follows: 0–1.0 min, 5% B; 1.0–3.0 min, 5–100% B; 3.0–3.5 min, 100% B; 3.5–4.0 min, 100–5% B; 4.0–6.5 min, 5% B. The flow rate was 0.25 mL/min, the column oven temperature was 60 °C, and the injection volume was 5 µL. Chromatographic separation was achieved using an ACQUITY UPLC BEH C18 column (50 × 2.1 mm, 1.7 µm particle size; Waters, USA). Detection was performed in positive ion mode using multiple reaction monitoring.

Since the clinically used i.v. formulation of TEIC comprises five major components (A_2-1_, A_2-2_, A_2-3_, A_2-4_, and A_2-5_), and one hydrolysis component (A_3-1_) (Fig. 1), the following ion transitions were monitored to detect each component: A_2-1_, from *m/z* 939.7 to 314.2; A_2-2_ and A_2-3_, from *m/z* 940.7 to 316.2; A_2-4_ and A_2-5_, from *m/z* 947.8 to 330.2; A_3-1_, from *m/z* 782.4 to 203.7 (9).

**Fig. 1.**
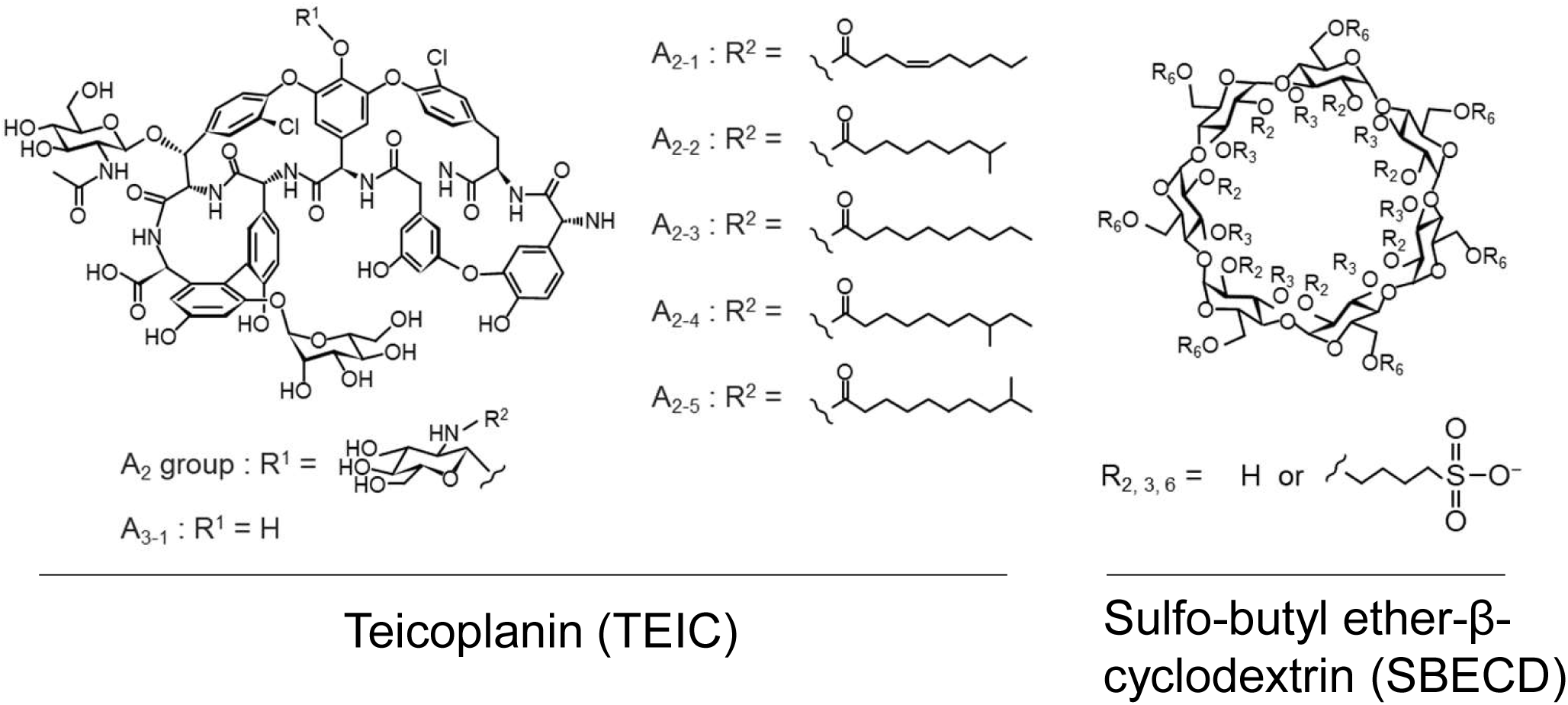
Chemical structures of teicoplanin (TEIC, left) and sulfo-butyl ether-β-cyclodextrin (SBECD, right).

The area under the plasma concentration–time curve (AUC) and the urinary excretion of TEIC were determined over 120 min after administration. The AUC and urinary excretion were calculated for total TEIC, and each individual component.

Statistical analyses were performed using GraphPad Prism version 8.0 software (GraphPad, San Diego, CA, USA). The data were analyzed using one-way analysis of variance; a *P*-value < 0.05 was considered statistically significant.

### Molecular docking simulation

Since TEIC A_2-2_ is the major component of the clinical formulation and its conformation is crystallographically characterized, only this dominant analog was considered for molecular modeling. The 3D structure of TEIC A_2-2_ was obtained from the Protein Data Bank (ID: 4PJZ) (10). The structure of SBECD was constructed using Avogadro software (ver. 1.2.0) (11) on the basis of the 3D structures of native β-CD that was retrieved from the Protein Data Bank (ID: 3CGT). Based on previous reports (12), the structure of SBECD was prepared by assuming that seven glucopyranose units were substituted with sulfobutyl ether, namely four at position C2 on the secondary face (residues 1, 3, 5, and 7) and three at position C6 on the primary face (residues 2, 4, and 6), in accordance with the average degree of substitution of the SBECD reagent (6.0-7.1). The molecular geometries of TEIC A_2-2_ and SBECD were fed into GAMESS (ver. 2023 R1) (13) for geometry optimization using the semi-empirical quantum chemistry method PM3. By treating the geometry-optimized structures of TEIC A_2-2_ and SBECD as ligands and receptors, respectively, molecular docking to obtain complexes of TEIC A_2-2_ with SBECD was performed with 100 runs using the Lamarckian genetic algorithm in AutoDock 4.2 (14). The receptor and ligand were assigned Gasteiger partial charges and the rotatable bonds were not defined. All other parameters were set to the default values for AutoDock 4.2.

### Effect of SBECD on TEIC protein binding in human serum

SBECD (40 mg/mL in distilled water) was diluted with human serum (Kojin Bio, Saitama, Japan) to prepare serum samples containing 0, 100, 300, and 1000 µg/mL of SBECD. These concentrations were within the range observed in clinical settings (15, 16). Aliquots of 380 µL of each serum sample were incubated at 37 °C for 30 min.

Subsequently, 20 µL of TEIC injection solution, prepared with distilled water to a concentration of 0.8 mg/mL, was added to achieve a final concentration of 40 µg/mL. This corresponded to the upper limit of the clinically targeted trough concentration of TEIC. The samples were then incubated at 37 °C for 60 min, with gentle mixing for 10 s every 10 min during incubation. After incubation, 300 µL of each sample was immediately collected and filtered through an ultrafiltration membrane (30 kDa, Millipore, Tokyo, Japan). TEIC calibration curves were generated using serum subjected to the same ultrafiltration procedure used for the matrix. The concentration of TEIC in the filtrate was quantified using the method described above, and the unbound fraction was calculated relative to the concentration of 40 µg/mL. Data were analyzed using one-way analysis of variance, and a P-value < 0.05 was considered statistically significant.

## Results

### Decreased TEIC plasma concentrations following i.v. PSCZ administration in patients who underwent HSCT

This study included eight patients who underwent HSCT and received both TEIC and PSCZ during the observation period. The plasma TEIC concentrations were measured under three conditions: without PSCZ administration, following oral PSCZ administration, and following i.v. PSCZ administration. Among these patients, TEIC was administered prior to PSCZ in five, while PSCZ was administered prior to TEIC in three. A total of 83 plasma concentration measurements were obtained: 23 without PSCZ administration, 27 following oral administration, and 33 following i.v. administration. The data points per person ranged from a minimum of five to a maximum of 18. The patient characteristics are summarized in Table 1. The average TEIC C/D ratio was 3.53 (standard deviation [SD], 1.29) in the absence of PSCZ, 3.10 (SD, 1.26) following oral PSCZ administration, and 2.23 (SD, 0.95) following i.v. administration (Fig. 2). A linear mixed-effects model demonstrated that the C/D ratio was significantly lower following i.v. PSCZ administration than without administration (estimate: -1.24; 95% confidence interval [CI], -1.74 to -0.74). After adjusting for serum albumin, eGFR, age, and sex as covariates, the reduction in the C/D ratio associated with i.v. PSCZ remained significant (estimate: -1.22; 95% CI, -1.84 to -0.60). This corresponds to a decrease of approximately 35% compared to the non-PSCZ condition.

**Fig. 2.**
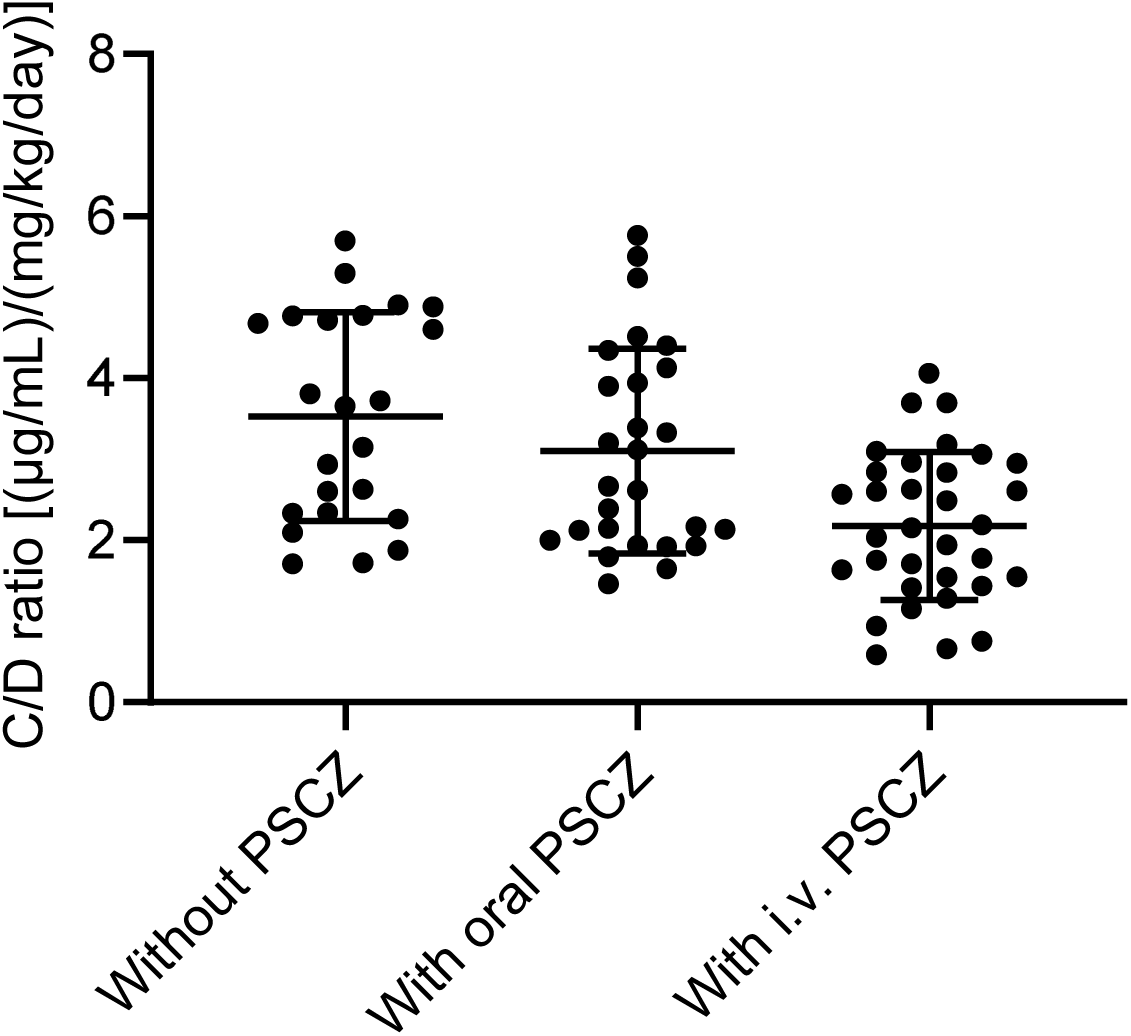
Trough plasma concentrations of TEIC in patients who underwent hematopoietic stem cell transplantation (HSCT) during periods without posaconazole (PSCZ) administration, following PSCZ oral administration, and following PSCZ intravenous administration. The TEIC concentration/dose (C/D) ratios were obtained from 83 plasma concentration measurements across eight patients: 23 who did not receive PSCZ, 27 who administered oral PSCZ, and 33 who received intravenous (i.v.) PSCZ. Bars indicate the mean C/D ratio, and error bars represent standard deviations.

**Table 1.** Characteristics of patients treated with teicoplanin and posaconazole.

|  |  |
| --- | --- |
| Dose of teicoplanin, mg/kg/day (mean ± S.D.) | 9.95 ± 3.64 |
| Male/female, n | 4/4 |
| Age, years (mean ± S.D.) | 45.9 ± 11.0 |
| Serum albumin, g/dL (mean ± S.D.) | 2.92 ± 0.45 |
| Estimated glomerular filtration rate, mL/min/1.73 m<sup>2</sup> (mean ± S.D.) | 101.5 ± 46.3 |
| Number of data points, n |  |
| Without posaconazole administration | 23 |
| Administered oral posaconazole | 27 |
| Administered intravenous posaconazole | 33 |

S.D., standard deviation. Posaconazole was administered at a dose of 300 mg/day regardless of body weight or route of administration.

In contrast, the effect of oral PSCZ administration on the TEIC C/D ratio relative to non-administration was not statistically significant (unadjusted: -0.36; 95% CI, -0.92 to 0.20; adjusted: -0.21; 95% CI, -0.88 to 0.45).

### Changes in TEIC pharmacokinetics following SBECD administration in rats

The above results suggest that the i.v. formulation of PSCZ is associated with reduced plasma TEIC concentrations. Because the dosage of PSCZ was identical regardless of the route of administration (300 mg/day), we focused on SBECD, which is present only in the i.v. formulation, as a potential factor influencing TEIC pharmacokinetics. To test this hypothesis, rats were pretreated with either an i.v. PSCZ formulation or SBECD followed by TEIC administration, and the AUC and urinary excretion of TEIC were evaluated (Fig. 3). Compared to rats pretreated with saline, those pretreated with either i.v. PSCZ or SBECD showed a significant decrease in TEIC AUC and a significant increase in urinary excretion (Fig. 3A–C). No significant differences were observed in the plasma concentrations immediately after administration (Figure 3A) or significant differences in urine volume (Fig. 3D) between the groups. Furthermore, the relative abundance of the individual components of the TEIC formulation, A_2-1_, A_2-2/2-3_, A_2-4/2-5_, and A_3-1_, was determined. Following the i.v. administration of PSCZ or SBECD, the AUC of the A_2_ group significantly decreased, whereas the urinary excretion significantly increased (Fig. 3E–G, 3I–K). In contrast, no significant effects were observed for A_3-1_ (Fig. 3H and 3L).

**Fig. 3.**
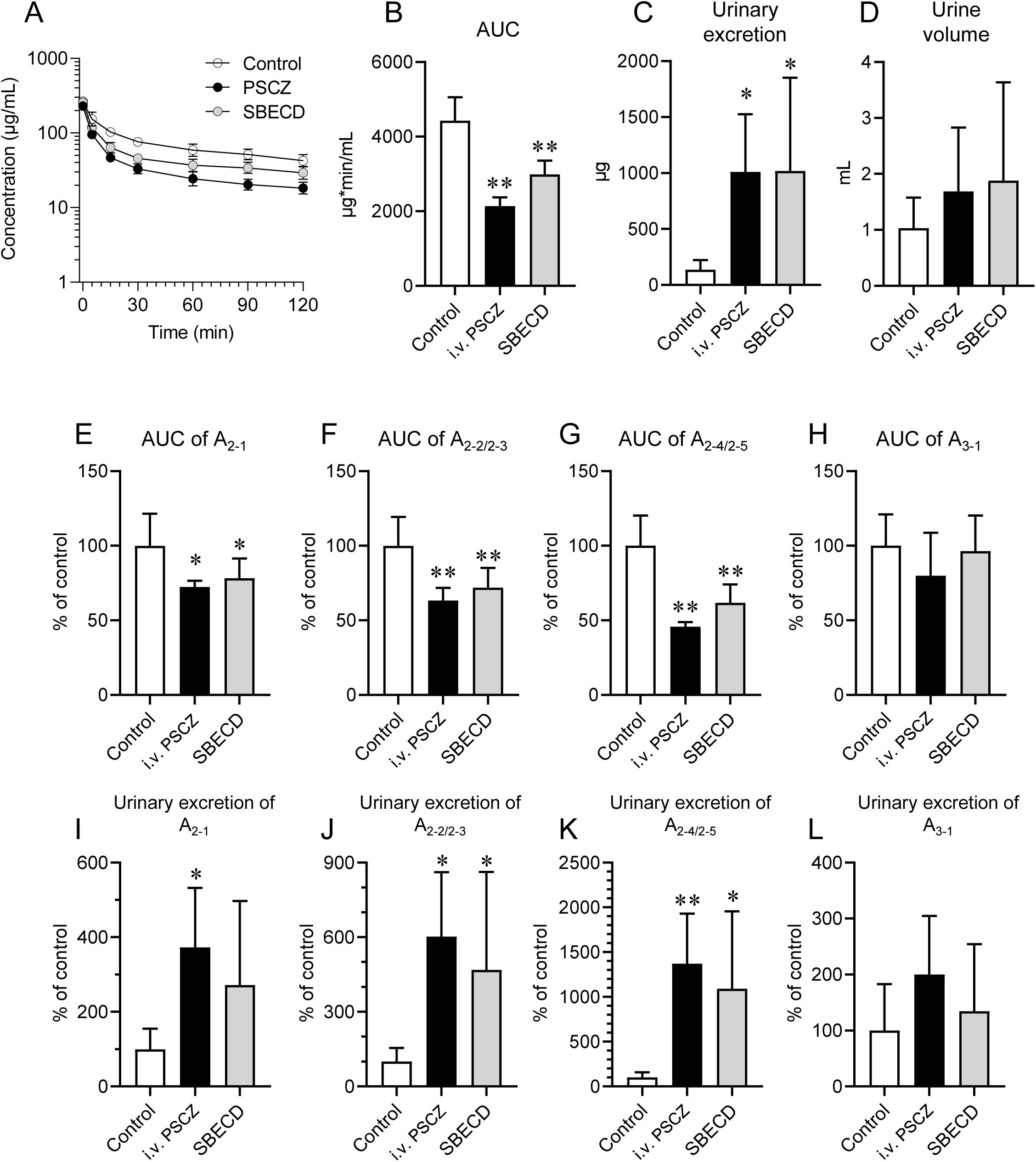
Pharmacokinetics of TEIC in rats. Male Wistar/ST rats (8 weeks old) received an intravenous injection via the femoral vein of saline (Control), PSCZ intravenous formulation (i.v. PSCZ), or SBECD alone (SBECD). Thirty minutes later, TEIC was administered intravenously. (A–D) The area under the plasma concentration–time curve (AUC) and urinary excretion of TEIC were evaluated over 120 min after TEIC administration. (E–L) The AUC and urinary excretion of individual TEIC components were evaluated and expressed as relative values to the Control group. \**P* < 0.05, \*\**P* < 0.01, significantly different from the Control group (n = 6 per group).

### Molecular modeling of TEIC A_2-2_ inclusion complex with SBECD

The abovementioned findings indicate that SBECD contributes to a reduction in TEIC plasma concentration, with a greater impact on the A_2_ group than A_3-1_.

Structural differences exist between these groups because only the A_2_ group possesses a long carbon chain (Fig. 1). Furthermore, a previous study showed that a β-cyclodextrin derivative, 2-hydroxypropyl-β-cyclodextrin (HP-β-CD), potentially prevents the hydrolysis of the glycosyl group with a hydrophobic side chain of dalbavancin, a semisynthetic glycopeptide with an analogous molecular scaffold of TEIC (17).

Therefore, we hypothesized that SBECD accommodates the side chain of TEIC, thereby inhibiting its binding to serum proteins. To test this postulation, we performed docking simulations for TEIC A_2-2_, a major analog of TEIC, against SBECD using AutoDock software. Through 100 docking runs, we obtained two probable binding modes for the TEIC A_2-2_ complex with SBECD: the dimethyl terminus of the TEIC A_2-2_ side chain was directed towards the primary face of the SBECD molecule for Type I (Fig. 4A), and the dimethyl terminus was directed towards the secondary face of the SBECD molecule for Type II (Fig. 4B). Regardless of the binding mode, most of the TEIC A_2-2_ hydrophobic side chain, which contributes to an increased protein-binding rate, was accommodated within the hydrophobic inner cavity of SBECD.

**Fig. 4.**
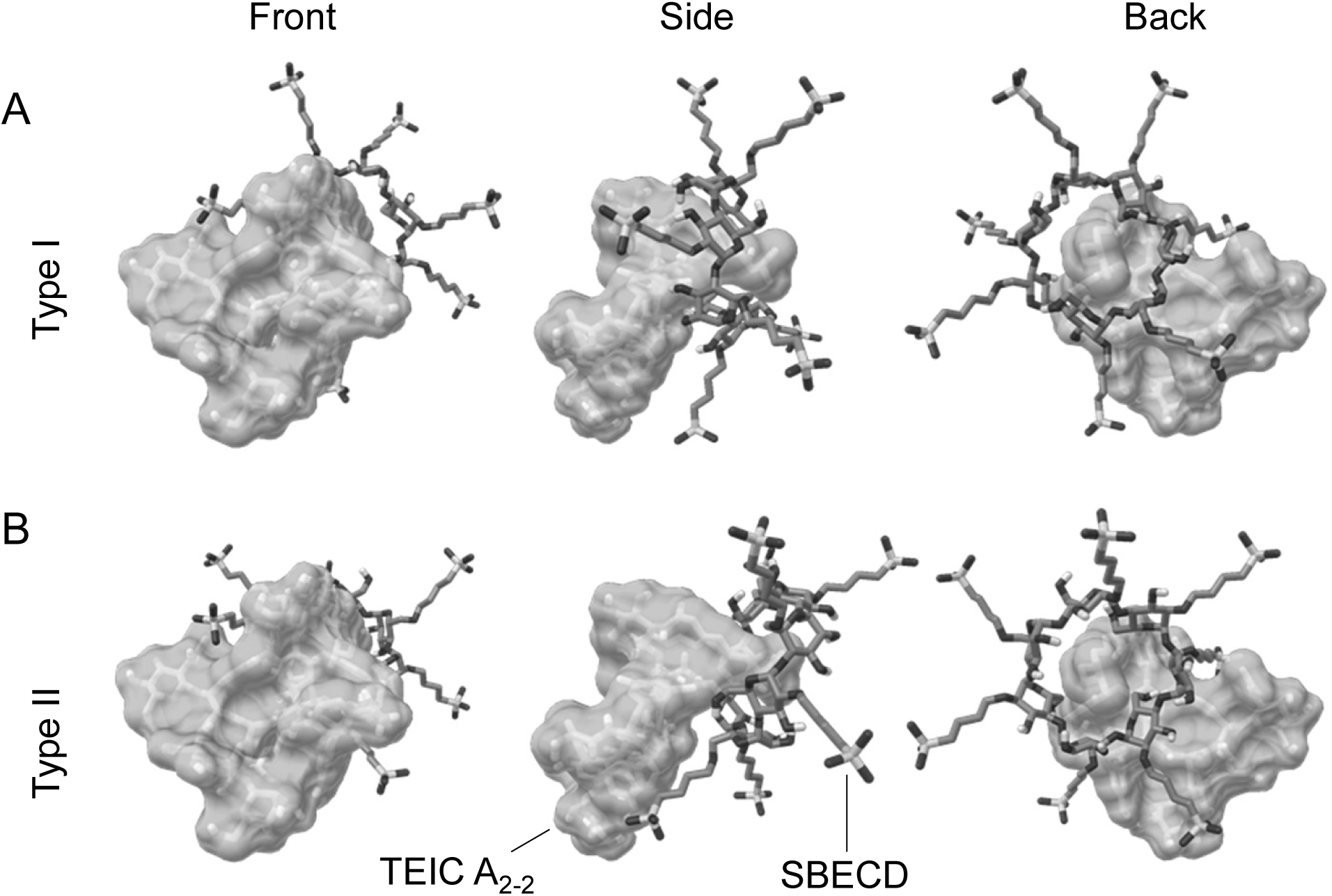
Estimated binding modes of TEIC A_2-2_ with SBECD. The lowest TEIC A_2-2_ binding energy conformations for SBECD in the most probable docking clusters from 100 runs. The dimethyl terminus of the TEIC A_2-2_ side chain is directed towards the primary face of the SBECD molecule in Type I (A) and is directed towards the secondary face of the SBECD molecule in Type II (B). The left, middle, and right panels show the front, side, and back views as the chemical structure of TEIC in Figure 1, respectively.

### Effect of SBECD on TEIC protein binding

The above findings suggest that SBECD exerts a strong influence on TEIC protein binding, particularly on the A_2_ group. Therefore, we evaluated the unbound fraction of TEIC in human serum containing clinically relevant concentrations of SBECD (Fig. 5). The results demonstrated that the unbound fractions of both the A_2_ group and A_3-1_ increased in a concentration-dependent manner with increasing SBECD concentration. Compared with the condition without SBECD, the unbound fractions of A_2-1_, A_2-2/2-3_, A_2-4/2-5_, and A_3-1_ increased by 3.4-fold, 9.6-fold, 25.9-fold, and 1.3-fold, respectively, at an SBECD concentration of 300 µg/mL.

**Fig. 5.**
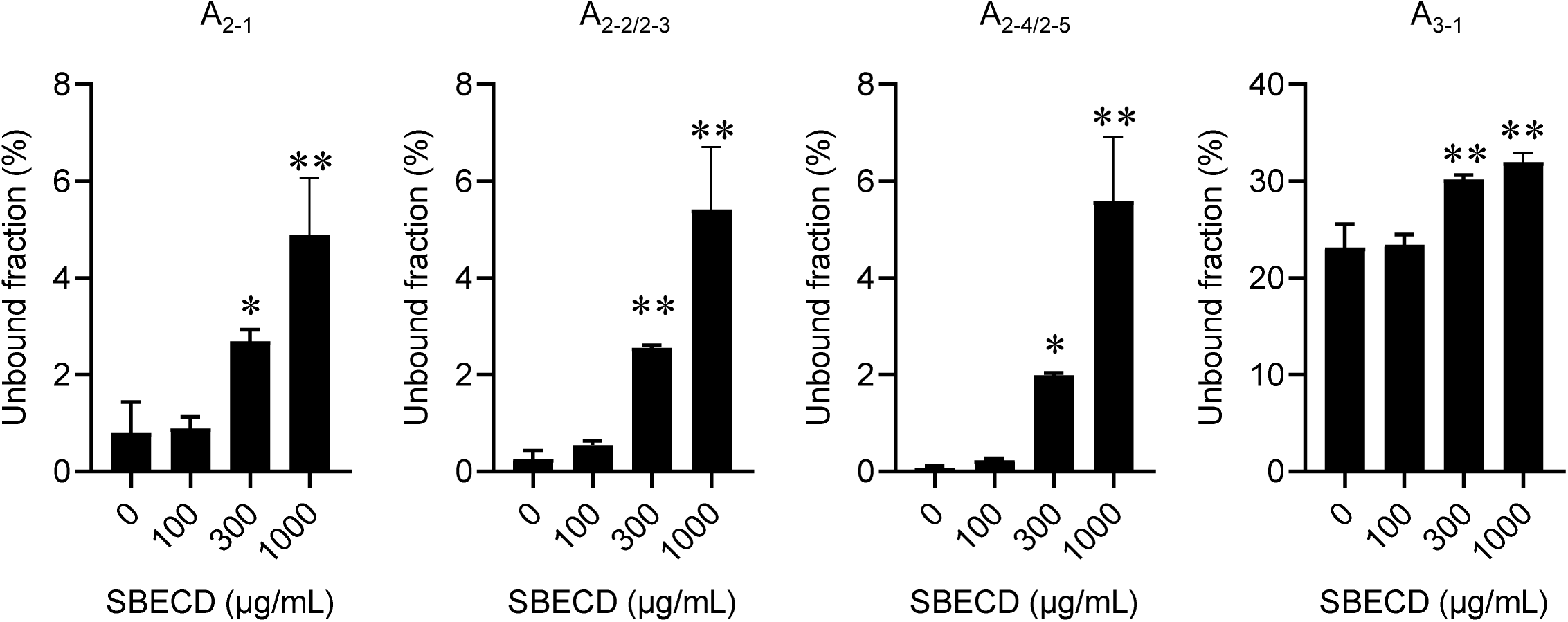
Unbound fraction of TEIC in human serum. Human serum containing various concentrations of SBECD was supplemented with TEIC. The unbound fraction of each TEIC component was determined. *P < 0.05, **P < 0.01, significantly different from the group without SBECD (“0”) (n = 3 per group).

## Discussion

Multiple antibiotics and antifungal agents are often used concurrently in patients undergoing HSCT, making it essential to fully understand the potential interactions between these drugs. In this study, we demonstrated, for the first time, that the co-administration of an i.v. formulation of PSCZ with TEIC in patients undergoing HSCT resulted in reduced TEIC plasma concentration. A key strength of this study is that the reproducibility of this phenomenon was confirmed by integrating clinical observations with an animal model. Moreover, by focusing on the pharmaceutical excipient SBECD, we further elucidated its impact on TEIC protein binding using in vitro assays and the mode of inclusion complex formation through molecular modeling.

SBECD is a derivative of β-cyclodextrin, composed of seven glucose units linked in a ring, with a hydrophobic inner cavity and a hydrophilic outer surface. This structure enables the encapsulation of hydrophobic drugs, thereby improving their aqueous solubility (18–20). Each vial of the i.v. PSCZ formulation (Noxafil® 300 mg) contains 6,680 mg of SBECD. In general, cyclodextrin–drug inclusion complexes are thought to have minimal effects on pharmacokinetics, as they dissociate rapidly after i.v. administration (21–27). However, exceptionally stable complexes are known to markedly alter drug elimination pathways. For instance, the rocuronium–sugammadex complex exhibits a high stability constant, whereas rocuronium alone is excreted through bile and rapidly eliminated in urine in the presence of sugammadex (21).

Similarly, drugs with an adamantane structure show particularly high affinity because of their close fit with the cyclodextrin cavity (28), and urinary excretion rates have been reported to increase from 0.1% to 37% in the presence of SBECD (29). In the current study, rats administered with either the i.v. PSCZ formulation or SBECD alone exhibited a significant increase in TEIC urinary excretion, supporting the interaction between TEIC and SBECD in vivo.

Semisynthetic lipoglycopeptides, such as the TEIC A_2_ group, telavancin, dalbavancin, and oritavancin, exhibit high protein binding and consequently much longer half-lives than vancomycin owing to their lipophilic side chains (30). Although the binding affinity of TEIC for human serum albumin (HSA), the predominant serum protein, has been reported (31) , its binding conformation remains unclear. Dalbavancin, which shares a large structural similarity with TEIC, binds to HSA by inserting its hydrocarbon chain into the hydrophobic pocket within subdomain IA of HSA, composed entirely of nonpolar amino acids (32). Given that TEIC, despite isoform differences, also possesses a long hydrocarbon chain and a conserved cyclic peptide core, it is likely to bind albumin in a similar manner. This hypothesis is consistent with our in vitro findings that showed a higher proportion of the unbound fraction of A_3-1_ than that of the A_2_ group. Furthermore, HP-β-CD has been shown to slowly hydrolyze the glycosidic linkage between the mannosyl aglycone and hydrophobic side chain of dalbavancin (17), suggesting that HP-β-CD interacts with and shields this region. The use of HP-β-CD as a pharmaceutical excipient to enhance the solubility of telavancin (Vibativ®) (33) and oritavancin (Kimyrsa®) (34) further supports our findings regarding TEIC–CD interactions. A_2-2_, A_2-3_, A_2-4_, and A_2-5_ are the major components of clinical TEIC formulations. In this study, SBECD significantly increased the unbound fraction of TEIC, particularly in the A_2_ group. These results suggest that the hydrophobic cavity of SBECD accommodates much of the hydrophobic side chain of TEIC A_2_, thereby reducing protein binding and leading to decreased plasma concentrations through enhanced renal excretion.

In patients who underwent HSCT, the C/D ratio of TEIC decreased by an average of 1.2 when the i.v. formulation of PSCZ was administered, compared with no administration. In other words, when TEIC was administered at 10 mg/kg/day, co-administration of i.v. PSCZ resulted in an approximate 12 µg/mL reduction in TEIC trough levels. However, this study had certain limitations. The clinical sample size in this study was limited, and large-scale prospective studies are needed to evaluate the magnitude of this effect more accurately. In addition, an assessment of how these plasma concentration changes affect the efficacy of TEIC should be conducted through pharmacokinetic-pharmacodynamic analyses. Moreover, since SBECD is widely used in injectable formulations beyond PSCZ, future studies should investigate potential interactions between TEIC and these SBECD-containing products, as well as with drugs possessing similar hydrophobic side chains.

Taken together, our findings suggest that the co-administration of i.v. PSCZ containing SBECD and TEIC may reduce the protein binding of the latter, thereby promoting its renal excretion. The findings of this study provide a new perspective for assessing the risk of drug–drug interactions when using SBECD-containing formulations.

## Acknowledgements

This work was supported by the grant from the Nakatomi Foundation. The funders had no role in study design, data collection and interpretation, or the decision to submit the work for publication.

## References

1. Roberts JA, Pea F, Lipman J. 2013. The clinical relevance of plasma protein binding changes. Clin Pharmacokinet 52:1–8.

2. Wilson AP. 2000. Clinical pharmacokinetics of teicoplanin. Clin Pharmacokinet 39:167–83.

3. Chambers HF, Kennedy S. 1990. Effects of dosage, peak and trough concentrations in serum, protein binding, and bactericidal rate on efficacy of teicoplanin in a rabbit model of endocarditis. Antimicrob Agents Chemother 34:510–4.

4. Hanai Y, Takahashi Y, Niwa T, Mayumi T, Hamada Y, Kimura T, Matsumoto K, Fujii S, Takesue Y. 2022. Clinical practice guidelines for therapeutic drug monitoring of teicoplanin: a consensus review by the Japanese Society of Chemotherapy and the Japanese Society of Therapeutic Drug Monitoring. J Antimicrob Chemother 77:869–879.

5. Lewis R, Niazi-Ali S, McIvor A, Kanj SS, Maertens J, Bassetti M, Levine D, Groll AH, Denning DW. 2024. Triazole antifungal drug interactions-practical considerations for excellent prescribing. J Antimicrob Chemother 79:1203–1217.

6. Lempers VJ, van den Heuvel JJ, Russel FG, Aarnoutse RE, Burger DM, Brüggemann RJ, Koenderink JB. 2016. Inhibitory Potential of Antifungal Drugs on ATP-Binding Cassette Transporters P-Glycoprotein, MRP1 to MRP5, BCRP, and BSEP. Antimicrob Agents Chemother 60:3372–9.

7. Soy D, López E, Ribas J. 2006. Teicoplanin population pharmacokinetic analysis in hospitalized patients. Ther Drug Monit 28:737–43.

8. Nair AB, Jacob S. 2016. A simple practice guide for dose conversion between animals and human. J Basic Clin Pharm 7:27–31.

9. Kim KY, Cho SH, Song YH, Nam MS, Kim CW. 2016. Direct injection LC-MS/MS method for the determination of teicoplanin in human plasma. J Chromatogr B Analyt Technol Biomed Life Sci 1008:125–131.

10. Berman HM, Westbrook J, Feng Z, Gilliland G, Bhat TN, Weissig H, Shindyalov IN, Bourne PE. 2000. The Protein Data Bank. Nucleic Acids Res 28:235–42.

11. Hanwell MD, Curtis DE, Lonie DC, Vandermeersch T, Zurek E, Hutchison GR. 2012. Avogadro: an advanced semantic chemical editor, visualization, and analysis platform. J Cheminform 4:17.

12. Rescifina A, Surdo E, Cardile V, Avola R, Eleonora Graziano AC, Stancanelli R, Tommasini S, Pistarà V, Ventura CA. 2019. Gemcitabine anticancer activity enhancement by water soluble celecoxib/sulfobutyl ether-β-cyclodextrin inclusion complex. Carbohydr Polym 206:792–800.

13. Barca GMJ, Bertoni C, Carrington L, Datta D, De Silva N, Deustua JE, Fedorov DG, Gour JR, Gunina AO, Guidez E, Harville T, Irle S, Ivanic J, Kowalski K, Leang SS, Li H, Li W, Lutz JJ, Magoulas I, Mato J, Mironov V, Nakata H, Pham BQ, Piecuch P, Poole D, Pruitt SR, Rendell AP, Roskop LB, Ruedenberg K, Sattasathuchana T, Schmidt MW, Shen J, Slipchenko L, Sosonkina M, Sundriyal V, Tiwari A, Galvez Vallejo JL, Westheimer B, Włoch M, Xu P, Zahariev F, Gordon MS. 2020. Recent developments in the general atomic and molecular electronic structure system. J Chem Phys 152:154102.

14. Morris GM, Huey R, Lindstrom W, Sanner MF, Belew RK, Goodsell DS, Olson AJ. 2009. AutoDock4 and AutoDockTools4: Automated docking with selective receptor flexibility. J Comput Chem 30:2785–91.

15. Luke DR, Tomaszewski K, Damle B, Schlamm HT. 2010. Review of the basic and clinical pharmacology of sulfobutylether-beta-cyclodextrin (SBECD). J Pharm Sci 99:3291–301.

16. Morris AA, Mueller SW, Rower JE, Washburn T, Kiser TH. 2015. Evaluation of Sulfobutylether-β-Cyclodextrin Exposure in a Critically Ill Patient Receiving Intravenous Posaconazole While Undergoing Continuous Venovenous Hemofiltration. Antimicrob Agents Chemother 59:6653–6.

17. Jakaria SM, Budil DE, Murtagh J. 2023. Strategies To Stabilize Dalbavancin in Aqueous Solutions; Section 3: The Effects of 2 Hydroxypropyl-β-Cyclodextrin and Phosphate Buffer with and without Divalent Metal Ions. Pharm Res 40:2027–2037.

18. Yan M, Wu S, Wang Y, Liang M, Wang M, Hu W, Yu G, Mao Z, Huang F, Zhou J. 2024. Recent Progress of Supramolecular Chemotherapy Based on Host-Guest Interactions. Adv Mater 36:e2304249.

19. Kali G, Haddadzadegan S, Bernkop-Schnürch A. 2024. Cyclodextrins and derivatives in drug delivery: New developments, relevant clinical trials, and advanced products. Carbohydr Polym 324:121500.

20. Crini G. 2014. Review: a history of cyclodextrins. Chem Rev 114:10940–75.

21. Loftsson T, Moya-Ortega MD, Alvarez-Lorenzo C, Concheiro A. 2016. Pharmacokinetics of cyclodextrins and drugs after oral and parenteral administration of drug/cyclodextrin complexes. J Pharm Pharmacol 68:544–55.

22. Kurkov SV, Loftsson T, Messner M, Madden D. 2010. Parenteral delivery of HPβCD: effects on drug-HSA binding. AAPS PharmSciTech 11:1152–8.

23. Piel G, Evrard B, Van Hees T, Delattre L. 1999. Comparison of the IV pharmacokinetics in sheep of miconazole-cyclodextrin solutions and a micellar solution. Int J Pharm 180:41–5.

24. Liu J, Qiu L, Gao J, Jin Y. 2006. Preparation, characterization and in vivo evaluation of formulation of baicalein with hydroxypropyl-beta-cyclodextrin. Int J Pharm 312:137–43.

25. Egan TD, Kern SE, Johnson KB, Pace NL. 2003. The pharmacokinetics and pharmacodynamics of propofol in a modified cyclodextrin formulation (Captisol) versus propofol in a lipid formulation (Diprivan): an electroencephalographic and hemodynamic study in a porcine model. Anesth Analg 97:72–9, table of contents.

26. Buggins TR, Dickinson PA, Taylor G. 2007. The effects of pharmaceutical excipients on drug disposition. Adv Drug Deliv Rev 59:1482–503.

27. Kurkov SV, Madden DE, Carr D, Loftsson T. 2012. The effect of parenterally administered cyclodextrins on the pharmacokinetics of coadministered drugs. J Pharm Sci 101:4402–8.

28. Leong NJ, Prankerd RJ, Shackleford DM, Mcintosh MP. 2015. The effect of intravenous sulfobutylether7 -β-cyclodextrin on the pharmacokinetics of a series of adamantane-containing compounds. J Pharm Sci 104:1492–8.

29. Charman SA, Perry CS, Chiu FC, McIntosh KA, Prankerd RJ, Charman WN. 2006. Alteration of the intravenous pharmacokinetics of a synthetic ozonide antimalarial in the presence of a modified cyclodextrin. J Pharm Sci 95:256–67.

30. Van Bambeke F, Van Laethem Y, Courvalin P, Tulkens PM. 2004. Glycopeptide antibiotics: from conventional molecules to new derivatives. Drugs 64:913–36.

31. Assandri A, Bernareggi A. 1987. Binding of teicoplanin to human serum albumin. Eur J Clin Pharmacol 33:191–5.

32. Ito S, Senoo A, Nagatoishi S, Ohue M, Yamamoto M, Tsumoto K, Wakui N. 2020. Structural Basis for the Binding Mechanism of Human Serum Albumin Complexed with Cyclic Peptide Dalbavancin. J Med Chem 63:14045–14053.

33. Wenzler E, Rodvold KA. 2015. Telavancin: the long and winding road from discovery to food and drug administration approvals and future directions. Clin Infect Dis 61 Suppl 2:S38–47.

34. Hoover RK, Krsak M, Molina KC, Shah K, Redell M. 2022. Kimyrsa, An Oritavancin-Containing Product: Clinical Study and Review of Properties. Open Forum Infect Dis 9:ofac090.

